# Mitigating reactive oxygen species production and increasing gel porosity improves lymphocyte motility and fibroblast spreading in photocrosslinked gelatin-thiol hydrogels

**DOI:** 10.1101/2024.01.14.574282

**Authors:** Tochukwu Ozulumba, Jonathan M. Zatorski, Abhinav Arneja, Jennifer H. Hammel, Thomas J. Braciale, Chance J. Luckey, Jennifer M. Munson, Rebecca R. Pompano

## Abstract

On-chip 3D culture systems that incorporate immune cells such as lymphocytes and stromal cells are needed to model immune organs in engineered systems such as organs-on-chip. Photocrosslinking is a useful tool for creating such immune-competent hydrogel cultures with spatial cell organization. However, loss of viability and motility in photocrosslinked gels can limit its utility, especially when working with fragile primary cells. We hypothesized that optimizing photoexposure-induced ROS production, hydrogel porosity or a combination of both factors was necessary to sustain cell viability and motility during culture in photocrosslinked gelatin-thiol (GelSH) hydrogels. Jurkat T cells, primary human CD4+ T cells and human lymphatic fibroblasts were selected as representative lymphoid immune cells to test this hypothesis. Direct exposure of these cells to 385 nm light and LAP photoinitiator dramatically increased ROS levels. Pretreatment with an antioxidant, ascorbic acid (AA), protected the cells from light + LAP-induced ROS and was non-toxic at optimized doses. Furthermore, scanning electron microscopy showed that native GelSH hydrogels had limited porosity, and that adding collagen to GelSH precursor before crosslinking markedly increased gel porosity. Next, we tested the impact of AA pretreatment and increasing gel porosity, alone or in combination, on cell viability and function in 3D GelSH hydrogel cultures. Increasing gel porosity, rather than AA pretreatment, was more critical for rescuing viability of Jurkat T cells and spreading of human lymphatic fibroblasts in GelSH-based gels, but both factors improved the motility of primary human CD4+ T cells. Increased porosity enabled formation of spatially organized co-cultures of primary human CD4+ T cells and human lymphatic fibroblasts in photo-crosslinked gels in a multi-lane microfluidic chip, towards modeling the lymphoid organ microenvironment. Some optimization is still needed to improve homogeneity between regions on the chip. These findings will enable researchers utilizing photocrosslinking methods to develop immunocompetent 3D culture models that support viability and function of sensitive lymphoid cells.

## INTRODUCTION

Research efforts focused on lymphoid bioengineering, i.e., the development of tissue-mimicking scaffolds to model lymphoid organ architecture and function, have witnessed increased attention in recent years.^1,2^ Such immune-competent tissue engineered scaffolds could lend themselves to mechanistic studies, disease modeling, drug development and personalized medicine amongst other applications. Hydrogels are often the scaffold of choice for tissue engineering applications like these due to their broad-ranging utility, elastic capacity, tunability and a network morphology similar to the extracellular matrices of tissues.^3^ However, developing hydrogel scaffolds that can adequately support models of the organized architecture of lymphoid organs and recapitulate key cellular functions remains an ongoing challenge for the bioengineering field.

Of the currently available methods to fabricate 3D hydrogel matrices, photocrosslinking is especially beneficial because it offers rapid and efficient polymerization, does not require elevated temperatures, provides user control over parameters such as gelation kinetics and physicochemical properties, allows fabrication of complex shapes and spatial distribution of cells, and enables better biomimicry of tissue structure.^4–6^ When applied in bioprinting, photocrosslinking offers higher spatial resolution, greater mechanical stability of 3D features and improved printing fidelity than other chemical crosslinking approaches.^3,7,8^ Similarly, cell-laden hydrogel constructs have been photopatterned with precise spatial and temporal control by lithography through a mask to mimic the complex architecture and local microenvironments of organs in vivo.^9–11^

While studies have reported successful culture of encapsulated cell lines^12–15^ and induced pluripotent stem cells^16–18^ within photocrosslinked hydrogels and 3D printed scaffolds, there is also evidence of photoreaction-induced damage to a host of cell types. Exposure to ultraviolet radiation, photoinitiator and the highly reactive free radicals generated during photocrosslinking could damage plasma membranes, DNA and proteins, causing harm to cells and impairing cell viability.^5,19,20^ Ultraviolet light has been associated with greater DNA damage than visible light.^20^ Also, high intensity irradiation decreased cell viability in photocrosslinked gels.^21–23^ In addition to radiation, photoinitiators exert variable effects on cell viability across different cell types,^24^ often with reduced cell viability in response to increasing photoinitiator concentration.^22,23,25–28^ The chemistry of the polymer itself also matters. Photocrosslinked hydrogels, developed using thiol-norbornene click chemistry,^10,12^ are considered more biocompatible than methacrylate-based hydrogels because they produce less reactive oxygen species (ROS)^29^ and support cell spreading.^30^ Yet, even thiol-norbornene chemistry can have delayed toxic effects on some cells.^26^

Apart from photoexposure-induced ROS production, scaffold properties such as pore size and porosity can affect cell behavior after encapsulation. Larger pore sizes, and the associated increase in porosity, was correlated to decreased tensile strength in 3D printed GelMA scaffolds.^15^ Stachowiak et al^31^ showed that porosity and potentially, interconnectedness of pores was critical for lymphocyte motility in hydrogel culture. Similarly, proliferation of human dermal fibroblasts was increased in 3D printed gelatin scaffolds with larger pores compared to scaffolds with smaller pores.^32^ Thus, achieving successful culture of cells in photocrosslinked gels requires careful optimization of macromer concentration, light intensity, exposure time and photoinitiator concentration to minimize risk of cell damage.

Previously, we showed that primary human T cells could be photopatterned in gelatin-thiol (GelSH)/PEG-norbornene (PEGNB) hydrogels with high viability at 12 hours.^10^ However, there was a significant decline in viability after 24-hour culture and lack of lymphocyte motility.^10^ We hypothesized that either ROS exposure, low porosity, or both must be optimized to enable multi-day culture of fragile immune cells (Figure 1). Therefore, towards the goal of patterning lymphoid tissue, here we systematically quantified photoexposure-induced ROS production and hydrogel porosity, and identified strategies to improve optimize multi-day viability and function of T cells and lymphatic fibroblasts after encapsulation in photocrosslinked hydrogels. We compared the impact of antioxidant pretreatment and hydrogel porosity on cell viability, fibroblast spreading, lymphocyte motility and activation-induced cytokine secretion in photocrosslinked GelSH/PEG-NB gels. Finally, as a proof-of-principle, we tested the ability of the system to allow spatial photopatterning of primary human T cells and lymphatic fibroblasts in a microfluidic chip.

**Figure 1.**
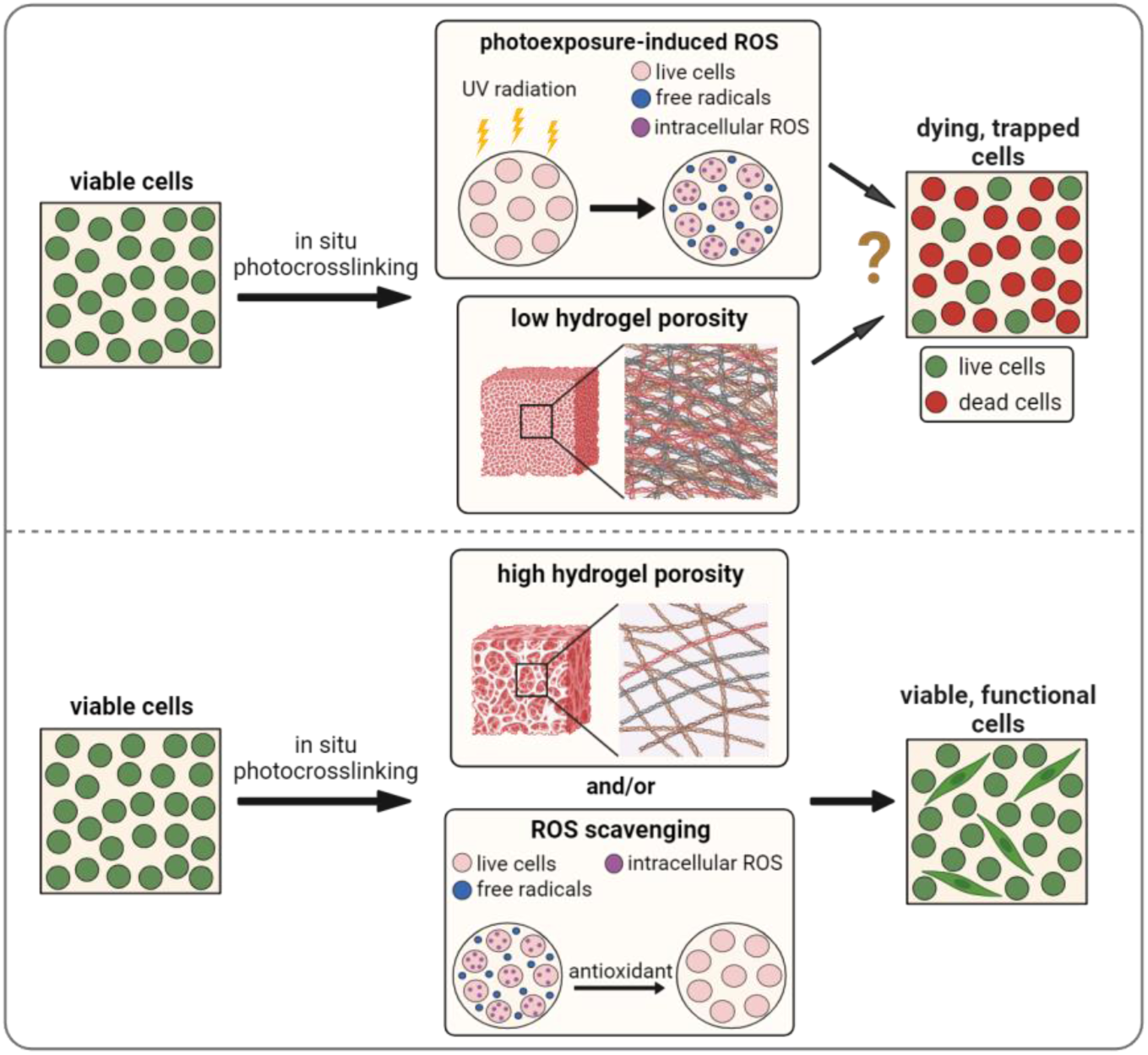
Schematic showing potential sources of toxicity to cells from in situ photocrosslinking (photoexposure-induced ROS and low hydrogel porosity). Figure created with BioRender.com.

## MATERIALS AND METHODS

### Cell sourcing and culture

Jurkat T lymphoblasts (Clone E6-1, TIB-152, ATCC, VA, USA) were obtained from Dr. Ken Hsu’s laboratory at the University of Virginia. Jurkat cells were cultured in RPMI media (Lonza) supplemented with 10% fetal bovine serum (FBS) (VWR, Lot No: 30121001), 2 mM L-glutamine (Gibco) and Penicillin (100 U/mL)/Streptomycin (100 µg/mL) (Pen/Strep) (Gibco). For culture, the cells were centrifuged at 200 *g* for 5 min, resuspended in 1 mL of media and counted using trypan blue exclusion.

Human lymphatic fibroblasts (HLFs) and the associated cell culture reagents (fibroblast media, trypsin/EDTA, trypsin neutralization solution, FBS, fibroblast growth supplement and Pen/Strep) were purchased from ScienCell Research Laboratories. 1x phosphate buffered saline (PBS) solution without calcium or magnesium was obtained from Lonza (MD, USA). HLFs were cultured in treated tissue culture flasks with fibroblast media supplemented with 2% FBS, Pen/Strep and fibroblast growth supplement. Cells were subcultured upon reaching 95% confluency according to the product sheet. Following cell treatment with trypsin/EDTA solution and subsequent detachment from the flask, the cells were centrifuged at 1000 rpm for 5 min, resuspended in 1 mL of media and counted using trypan blue exclusion. New flasks were set up at a seeding density of 5000 cells/cm^2^ and passaged again at 95% confluency.

Human naive CD4+ T cells were purified from platelet apheresis collars obtained from healthy donors, using the EasySep™ Naïve CD4+ T cell isolation kit (STEMCELL Technologies) according to isolation procedures reported previously.^10^ Cells were cultured in serum-free AIM V media (Gibco) supplemented with 10 ng/mL recombinant human IL-7 (R&D Systems). For culture, the cells were centrifuged at 400 *g* for 5 min, resuspended in 1 mL of media and counted using trypan blue exclusion.

### ROS quantification upon cell stimulation with H_2_O_2_ or light and/or LAP and impact of antioxidant pretreatment

The effect of photocrosslinking conditions and reagents on cellular ROS levels was assessed using a 2′,7′-Dichlorodihydrofluorescein diacetate (DCFH-DA) assay protocol modified from literature.^33^ Following cell uptake of DCFH-DA, intracellular esterases cleave the acetate groups yielding dichlorodihydrofluorescein (DCFH). Oxidation of DCFH generates the fluorescent DCF which has an excitation/emission spectra of 485/535 nm.^34^ The DCFH-DA reagent was purchased from Sigma Aldrich and stored at –20 °C. A 5 mM stock solution was prepared by dissolving DCFH-DA in DMSO at room temperature, and single-use aliquots were stored at –20 °C. The DCFH-DA stock solution was diluted 100-fold in serum-free media to obtain the 50 µM working solution. Next, cells were labeled with the DCFH-DA probe by resuspending the cell pellet in 2 mL of 50 µM DCFH-DA solution in serum-free media and incubating at 37 °C for 40 min. After incubation, the cell suspension was centrifuged to remove the DCFH-DA solution (Jurkat cells – 200 *g* for 5 min, HLFs – 1000 rpm for 5 min and primary CD4+ T cells – 400 *g* for 5 min). Next, cells were washed twice with 3 mL of 1x PBS by centrifugation as above to remove excess probe. DCFH-DA labeled Jurkat cells, naive CD4+ T cells and HLFs were seeded in 96-well plates in 100 µL volumes at densities of 1 × 10^5^, 1 × 10^6^ and 2 x 10^4^ cells/well respectively. Black-walled well plates were used for the cell stimulation experiments to limit well-to-well fluorescence crosstalk. HLFs were specifically incubated for 24 h to allow for cell adhesion and >80% confluency in the plate wells before conducting any experiments. Seeded cells were treated with LAP (0.05%, 0.1% v/v), exposed to light (12.5, 25, 50 mW/cm^2^) for 10 sec, a combination of light (25 mW/cm^2^) and 0.05% LAP for 10 sec, or H_2_O_2_ (500 µM). After 24-hour culture, DCF fluorescence intensity was measured with a Biotek plate reader at excitation and emission wavelengths of 485 nm and 533 nm respectively.

For cell pretreatment with ascorbic acid (AA), a 50 mM AA stock solution was prepared by dissolving AA (Sigma Aldrich) in the relevant complete culture media. Cells were resuspended at 2 – 4 x 10^6^ cells/mL in a 2 mL AA solution contained in a 50 mL tube (varying concentrations of 1.56 – 25 mM AA in media) and incubated at 37 °C for 5 min. Afterwards, the cell suspension was centrifuged and the cell pellet resuspended in complete culture media to a concentration of 1 x 10^6^ cells/mL in a 50 mL tube. For cell pretreatment with N-acetylcysteine (NAC), an 80 mM NAC stock solution was prepared by dissolving NAC (Sigma Aldrich) in the relevant complete culture media. Jurkat cells were seeded at 1 x 10^5^ cells/mL in a 0.1 mL solution of NAC (varying concentrations of 1.25 – 40 mM) in a 96-well plate and incubated at 37 °C for 24 h.

### Cell viability studies

Jurkat cells, naive CD4+ T cells and HLFs were seeded in 96-well plates at densities of 1 × 10^5^, 1 × 10^6^ and 2 x 10^4^ cells/well respectively. Cells were stimulated with LAP, light, or light + LAP as above, and cell viability was measured after 24-hour incubation using the lactate dehydrogenase (LDH), 3-(4,5-dimethylthiazol-2-yl)-5-(3-carboxymethoxyphenyl)-2-(4-sulfophenyl)-2H-tetrazolium, inner salt (MTS) and Live/Dead assays. The MTS assay measures metabolic activity of cells on the principle that viable cells reduce tetrazolium compounds to generate UV-active formazan products.^35^ On the other hand, the LDH assay measures the levels of LDH, an intracellular enzyme, as an indicator of membrane damage.^36^ MTS and LDH assay kits were both purchased from Promega. For the MTS assay, spent cell culture media was removed from the wells and 120 µL of reconstituted MTS reagent was added to each well. The plate was incubated at 37°C, 5% CO_2_ for 2 h after which MTS absorbance was measured at 490 nm using a plate reader (BMG Labtech Clariostar). For the LDH assay, 50 µL of cell culture media was transferred to a new plate and 50 µL of the substrate mix was added. The plate was incubated at room temperature in the dark for 30 min after which the reaction was stopped by adding 50 µL of stop solution. LDH absorbance was measured at 492 nm using a plate reader. Both assays included untreated cells and cells lysed with Triton X (0.8% (v/v)) for 30 min as experimental controls.

For the Live/Dead assay, calcein AM and 4′,6-diamidino-2-phenylindole (DAPI) were purchased from Thermo Fisher Scientific. Killed cell controls were prepared by incubating untreated cells with 35% ethanol (Thermo Fisher Scientific) at 37 °C for 10 min. After 24-hour incubation of stimulated cells, the culture plate was centrifuged at 200 *g* (Jurkat cells) or 400 *g* (naive CD4+ T cells) for 5 min. Spent media was removed from the wells and the cell pellets were rinsed 2 times with 100 µL of PBS per well. Cells were resuspended in a staining solution of calcein AM (5 µM) and DAPI (1 µM) in 1x PBS at 100 µL per well and the plate was incubated at 37 °C for 30 min. Afterwards, the plate was centrifuged to remove the stain solution and the cell pellet was rinsed with 100 µL of 1x PBS per well. Cells were resuspended in 100 µL of 1x PBS and imaged using an upright Zeiss AxioZoom widefield microscope. Microscope and filter information are provided in the Supplementary Information. Fluorescence images were acquired with Zeiss Zen 3 software and analyzed using ImageJ (v1.53t).

### Hydrogel preparation and characterization

#### Collagen/fibrinogen (Col/Fib) gels

Rat tail collagen I (5 mg/mL) and fibrinogen were purchased from Ibidi and Sigma Aldrich respectively. 10x PBS, distilled H_2_O, NaHCO_3_ and 1x PBS were all obtained from Gibco. Fibrinogen was dissolved in 1x PBS to a concentration of 6 mg/mL, with rotation on an orbital shaker. Collagen/fibrinogen hydrogels were prepared following a modification of the manufacturer’s protocol.^37^ After incubation on ice for 10 min, 20 µL of 10x PBS, 6 µL of 1M NaOH, 49 µL of H2O, 5 µL of 7.5% NaHCO_3_, 50 µL of 1x PBS, 120 µL of collagen I and 50 µL of fibrinogen (6 mg/mL) were sequentially combined, with mixing after each step to ensure sample homogeneity. Final concentrations of collagen and fibrinogen in the precursor solution were 2 mg/mL and 1 mg/mL respectively. The precursor solution was incubated at 37 °C for 30 min to induce gelation.

#### Photocrosslinked gelatin-thiol (GelSH) gels

Thiol-modified gelatin (GelSH) and lithium phenyl-2,4,6-trimethylbenzoyl phosphinate (LAP) were purchased from Sigma Aldrich, while norbornene-functionalized polyethylene glycol (PEG-NB; 8-arm 20 kDa) was obtained from Jenkem Technologies. Photocrosslinked GelSH hydrogels were prepared following a modified procedure from Ortiz-Cárdenas et al.^10^ Gel precursor was prepared by combining 5% GelSH (w/v), 0.05% (w/v) LAP and 10 mM PEG-NB in 1x PBS. Photo-crosslinking was performed at a wavelength of 385 nm, light intensity of 25 mW/cm^2^ and exposure time of 10 sec, unless otherwise indicated. GelSH/collagen gels were prepared by combining GelSH/PEGNB and collagen gel precursor solutions in defined volume ratios in no special order, followed by pipette mixing and photocrosslinking as above. After crosslinking, the GelSH/collagen gels were incubated at 37 °C for 30 min followed by addition of media.

### Scanning electron microscopy of hydrogels

Gel precursor was loaded into an open-ended plastic cylinder (200 µL/cylinder) and either photocrosslinked (for GelSH), incubated at 37 °C for 30 min (for collagen/fibrinogen) or a sequential combination of both procedures (for GelSH/collagen). The gels were then flash frozen in liquid nitrogen and lyophilized for 24 h. Freeze dried gel sections were mounted on aluminum specimen stubs with double-sided carbon tape and coated with platinum (24 nm thickness) using a Cressington 108 Auto/SE sputter coater. Coating was run under argon flow at 30 mA and 0.05 mbar for 60 sec. Images were acquired with a FEI Quanta 650 scanning electron microscope at an accelerating voltage of 5 kV.

### Microfluidic device fabrication

One-layer microfluidic devices were fabricated using standard soft lithography methods as detailed elsewhere.^10^ PDMS layers and 1 mm thick glass slides were ozone treated for 10 sec, bonded together and incubated at 120 °C for 10 min. Bonded chips were covered with tape to limit dust contamination and stored at room temperature. Two versions of the chip were used, taking advantage of chips already available in our laboratory. Chip schematics are provided in the Supplementary Information.

For culturing Jurkat cells, the chip had four parallel channels separated by arrays of hexagonal micropillars (100 µm in diameter, with 50 µm spacing between pillars). The outer channels each had 8-mm diameter inlets and outlets that served as media reservoirs, while the inner gel channels had 0.75-mm inlets and outlets. Excluding the converging, angled channel regions, the media lanes were 2.45 x 0.13 x 14.81 mm^3^ (W x H x L) and the gel lanes were 1.4 x 0.13 x 15.4 mm^3^. For culturing primary CD4+ T cells, we used a chip with smaller dimensions to ensure that added stimuli quickly diffused throughout the width of the gel. In this ‘smaller’ chip, 5 parallel channels were separated by arrays of hexagonal micropillars (100 µm in diameter, with 50 µm spacing between pillars). The inlet/outlet diameters of the outer media channels and inner gel channels were 8 and 0.75 mm respectively. Excluding the converging, angled channel regions, the media lanes were 2.05 x 0.13 x 3.5 mm^3^ and the gel lanes were 0.3 x 0.13 x3.5 mm^3^.

### Cell encapsulation and analysis in hydrogels

Cell-laden gel was prepared by centrifuging cells and resuspending the pellet in the gel precursor before thermal incubation (Col/Fib) or photocrosslinking (GelSH and GelSH/collagen). For HLF culture, 50 µL of cell-laden gel (3 x 10^5^ cells/mL) was dispensed into a 96-well plate. For on-chip experiments with Jurkat cells, 10 µL of cell-laden precursor (1 x 10^6^ cells/mL) was loaded into the center lane of the larger chip. For primary CD4+ T cells, 2.5 µL of cell-laden gel (1 x 10^6^ cells/mL) was loaded into the smaller chip. After gelation, 150 µL of media was added to each of 4 chip reservoirs (total volume of 600 µL) or plate well (100 µL) for multi-day culture. In some experiments, CD4+ T cells were activated on-chip by adding ImmunoCult™ Human CD3/CD28 T Cell Activator (StemCell Technologies) to the culture media. Conditioned culture media was collected from the chip reservoirs after 72-hour culture and stored at –20 °C. Secreted IFN**γ** concentrations in the conditioned culture media were measured using an ELISA kit (BioLegend).

For Live/Dead staining of 3D cultures in the microfluidic device, 100 µL of calcein AM/DAPI stain solution was added to 2 homolateral chip reservoirs, and the chip was incubated at 37 °C for 2 h. During this time, the staining solution visibly diffused through the cell chamber to the opposite reservoir. Fluorescence images were acquired with an AxioZoom widefield microscope and images were analyzed using ImageJ.

For motility experiments, naive primary human CD4+ T cells were cultured on-chip for 24 h with media spiked with 1.22 ng/mL of recombinant human CCL21 (Peprotech). Brightfield images were acquired using transmitted light in time series mode (1 image was collected every 30 sec for 10 cycles) with an AxioZoom microscope. Individual cells (15 – 30) were tracked for image analysis using Cell Tracker (v1.1).^38^

### Fibroblast and T cell co-culture

To set up a co-culture of HLFs and CD4+ T cells on-chip, each cell-laden gel precursor was prepared separately and sequentially photopatterned following the respective protocol for each gel formulation. HLF-laden precursor was crosslinked in the center lane of the chip (smaller chip) at a seeding density of 5 x 10^6^ cells/mL. Precursor containing naïve CD4+ T cells (10 x 10^6^ cells/mL) was crosslinked in the 2 outer lanes of the chip. After gelation, 150 µL of media was added to each of 4 chip reservoirs (total volume of 600 µL) for 2-day culture. CD4+ T cells were activated on-chip by adding ImmunoCult™ Human CD3/CD28 T Cell Activator to culture media. To visualize inter-channel cell migration, 300 µL of a staining solution containing PE-conjugated anti-CD4 antibody (0.14 µM) and Calcein AM (5 µM) was added to two homolateral chip reservoirs and the chip was incubated at 37 °C for 2 h. The chips were rinsed twice by replacing the stain solution with 300 µL PBS and incubating at 37 °C for 30 min, then repeating. Fluorescence images were acquired using an inverted AxioZoom widefield microscope and representative images were prepared using ImageJ.

### Statistical analysis

Statistical tests and curve fits were performed using GraphPad Prism 10.0.1. Where indicated, a 4-parameter log(inhibitor) vs response curve was used, defined as *Y = Bottom + (Top-Bottom)/(1+10^((LogIC50 – X)*HillSlope))*. All bars indicate mean and standard deviation unless otherwise noted.

## RESULTS AND DISCUSSION

### Exposure to light and LAP induces synergistic ROS production in Jurkat T cells

We selected the thiol-norbornene based GelSH/PEG-NB for this work because it generates less ROS than methacrylate gels.^29^ LAP was chosen as the photoinitiator because it offers efficient polymerization at both ultraviolet (365 nm) and visible light (405 nm), has greater aqueous solubility and supports viability of encapsulated cells.^39^ Cell-laden gel precursor was photocrosslinked at 385 nm to leverage the rapid polymerization offered by shorter wavelengths while limiting cell damage.

To compare the impact of ROS production during photocrosslinking and hydrogel porosity on viability and motility of encapsulated cells, we first characterized each parameter, starting with ROS production and its effect on viability. We chose Jurkat cells for these initial screening experiments as they are a readily available model of T cells that is not subject to variation between donors, although as a proliferative cell line they may be expected to be more hardy than primary T cells. To quantify intracellular ROS levels, we used the fluorescence-based DCFH-DA assay (Figure 2b). A calibration curve measuring DCF fluorescence intensity in Jurkat cells increased linearly with the concentration of H_2_O_2_ in the media (Figure 2c), as expected.^40–43^

**Figure 2.**
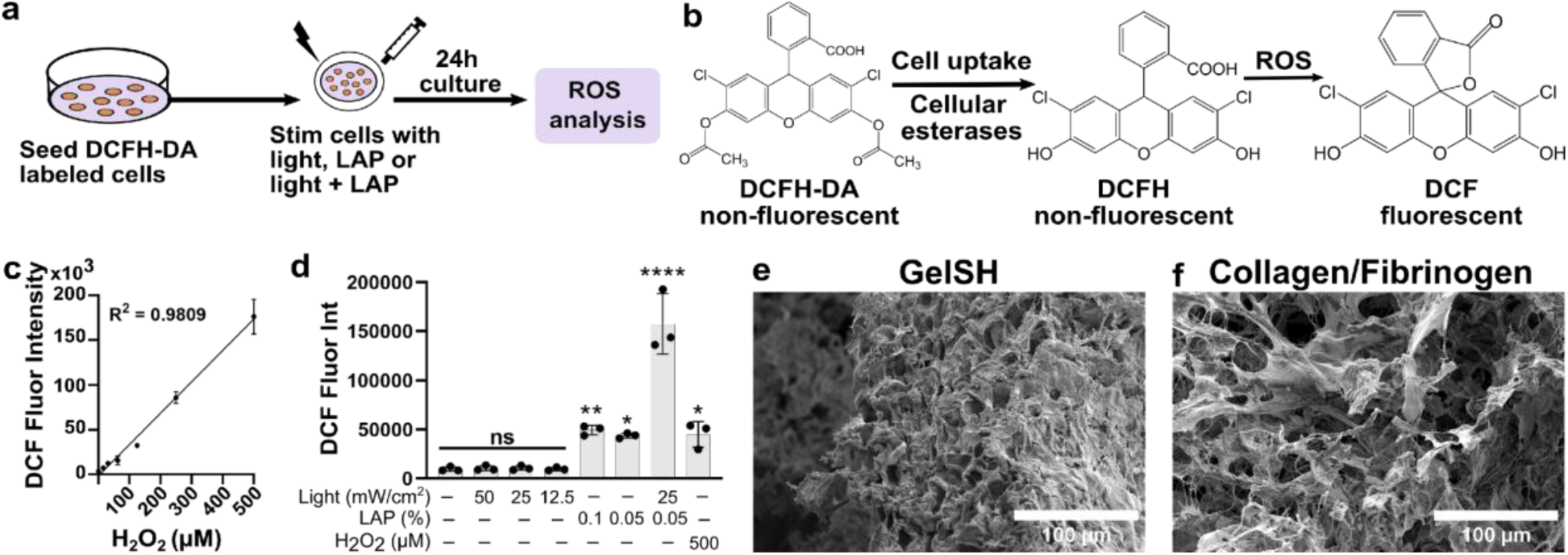
Characterization of ROS production and low porosity in photocrosslinked gelatin-thiol/PEG-norbornene hydrogels. (a) Schematic of the experimental procedure for measurement of ROS production in cells. (b) Principle of the DCFH-DA assay. (c) Standard curve from the DCFH-DA assay, plotting DCF fluorescence intensity in Jurkat cells versus H_2_O_2_ concentration added to the media. The black line represents the line of best fit (R^2^ = 0.9809), while symbols and error bars show mean and standard deviation. (d) ROS assay results showing DCF fluorescence intensities in Jurkat cells 24 h after exposure to 385 nm light at the indicated intensities (50, 25 and 12.5 mW/cm^2^), LAP, light + LAP or 500 μM H_2_O_2_ (n = 3). Bars show the mean with standard deviation; each dot is one biological replicate. One-way ANOVA with Dunnett’s multiple comparisons, ns p > 0.05, *p < 0.05, **p < 0.01, ****p < 0.0001 versus untreated cells. (e, f) SEM images of GelSH (e) and Col/Fib (f) hydrogels showing differences in gel porosity. Scale bar = 100 μm.

To quantify ROS levels in lymphocytes after photoexposure,^10^ we exposed Jurkat cells to 385 nm light, intensities of 12 – 50 mW/cm^2^ and LAP concentrations of 0.05 – 0.1% (w/v) in the absence of gel components. Cellular ROS levels did not increase in response to 385 nm light alone, similar to the observation made by Ruskowitz and DeForest.^44^ In contrast, ROS levels increased after 24-hour culture with LAP alone (4.5-fold and 5.1-fold over untreated cells for 0.05% LAP and 0.1% LAP respectively), similar to treatment with 500 µM H_2_O_2_, and they increased dramatically and synergistically (16.3-fold over untreated cells) following exposure to the combination of light and LAP (Figure 2d). A similar trend of increased potency of light + LAP compared to LAP was reported by Nguyen et al.^28^ Next, we tested the impact of exposure to light and/or LAP on Jurkat cell viability using the LDH, MTS and Live/Dead staining assays. Interestingly, although LDH absorbance was similar to the untreated cells in all test groups (Figure S1a), MTS absorbance in the light + LAP group was significantly decreased (1.7-fold) compared to untreated cells, matching ROS data (Figure S1b). This suggested that while Jurkat cell exposure to light + LAP had minimal impact on membrane integrity, metabolic activity was significantly impaired.

### Scanning electron microscopy reveals low porosity of GelSH gels

As prior reports indicated that gel porosity was critical to lymphocyte and fibroblast function in 3D culture,^31,45^ we characterized the morphology of the GelSH/PEGNB hydrogel using scanning electron microscopy (SEM). Thermally setting collagen-based hydrogels are routinely used for 3D cell culture^46–49^ and support lymphocyte migration,^50,51^ so they were included in this study as a positive control for optimal cell viability and function. SEM images of GelSH gels (Figure 2e) revealed a densely packed morphology in sharp contrast to the more porous network of collagen/fibrinogen (Col/Fib) gels (Figure 2f). Therefore, next we tested approaches both to mitigate photocrosslinking-induced ROS production and to increase the porosity of GelSH hydrogels.

### Pretreatment with antioxidants mitigates light + LAP-induced intracellular ROS production

We hypothesized that pretreating cells with an antioxidant would mitigate the ROS production induced by light + LAP, and we selected two antioxidants, NAC and AA to test this hypothesis. NAC, the acetylated variant of the amino acid L-cysteine, is used in clinical settings for treating diseases characterized by chemical toxicity or free oxygen radical production such as polycystic ovary syndrome, acetaminophen overdose and heavy metal poisoning, with minimal side effects.^52,53^ The antioxidant property of NAC is mediated via scavenging free radicals, reducing disulfide bonds in proteins and serving as a precursor for synthesis of glutathione,^52,53^ a critical antioxidant.^54^ On the other hand, AA (vitamin C) acts as an antioxidant by scavenging free radicals, stopping free radical chain reactions and reducing oxidized glutathione.^55,56^ AA has been shown to prevent lipid peroxidation and mitigate exercise-induced free radical production.^57^ Following overnight (24 h) pretreatment with these molecules, both NAC and AA mitigated the ROS induced by 500 µM H_2_O_2_ treatment in Jurkat cells (Figure S2). However, AA was more effective than NAC at reducing the comparatively larger quantity of ROS produced by light + LAP (385 nm light, 25 mW/cm^2^, 0.05% w/v LAP), particularly at lower doses (Figure 3b, c). Whereas NAC had an IC50 value of 940 mM, AA was more than 1000-fold lower at 0.44 mM. Therefore, AA was selected for further testing.

**Figure 3.**
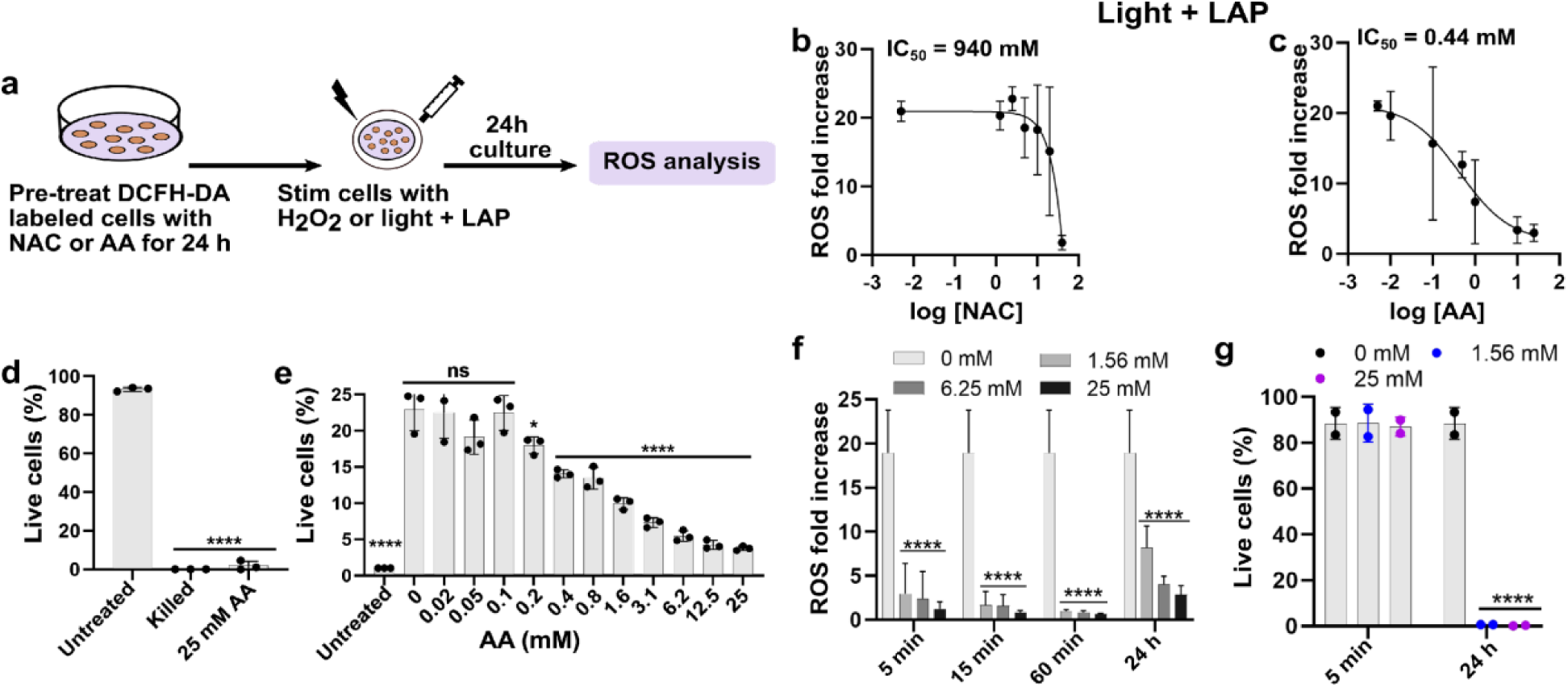
AA was more effective than NAC at mitigating light + LAP-induced ROS in Jurkat cells. (a) Schematic of the experimental procedure. (b, c) ROS fold increase in Jurkat cells after overnight (24 h) antioxidant pretreatment with NAC (1.25 – 80 mM) (b) or AA (1.56 – 25 mM) (c), followed by exposure to 385 nm light (25 mW/cm^2^) + LAP (0.05% (w/v), and another 24-hour culture (n = 3). Values were fit with a 4-parameter log(inhibitor) vs response curve. (d) Jurkat cell viability after overnight treatment with 25 mM AA (n = 3), in the absence of light or LAP exposure. One-way ANOVA with Dunnett’s multiple comparisons, ****p < 0.0001 versus untreated cells. (e) AA dose-response curve showing ROS fold increase in Jurkat cells after overnight pretreatment with varying doses of AA (0.02 – 25 mM) (n = 3), followed by light + LAP exposure. One-way ANOVA with Dunnett’s multiple comparisons, ns p > 0.05, *p < 0.05, ****p < 0.0001 versus no AA control (0 mM). (f) ROS fold increase in Jurkat cells as a function of AA dose and pretreatment time prior to light + LAP exposure (n = 4). Two-way ANOVA with Dunnett’s multiple comparisons, ****p < 0.0001 versus no AA control (0 mM). (g) Jurkat cell viability after AA pretreatment (1.56 mM vs 25 mM) for 5 min and 24 h without light + LAP exposure (n = 3). Two-way ANOVA with Dunnett’s multiple comparisons, ****p < 0.0001 versus no AA control (0 mM). Each dot indicates an independent experimental repeat.

### Optimizing AA dose and pretreatment time to maximize efficacy and cytocompatibility

T cells are sensitive to redox state, and optimal cell function hinges on a balance between pro-oxidant and antioxidant factors. Low (non-zero) ROS levels are critical for T cell survival, while increased ROS buildup during oxidative stress can result in apoptosis or necrosis.^58^ Therefore, we tested the effect of high concentrations of antioxidant on cell viability, even in the absence of photoexposure. Although overnight pretreatment with AA (25 mM) significantly mitigated light + LAP-induced ROS in Jurkat cells (Figure 3c), live/dead staining of cells pretreated overnight with 25 mM AA revealed that AA was itself toxic to the cells and resulted in cell death at that dose and treatment time (Figure 3d). Thus, optimization of AA pretreatment dose and duration was needed to avoid direct AA toxicity to the cells.

Titrating the AA dose used for overnight pretreatment prior to light + LAP exposure indicated significant mitigation of light + LAP-induced ROS at concentrations as low as 0.2 mM, with a 50% reduction at 0.75 mM (IC50 value) (Figure 3e). Hypothesizing that shorter pretreatment times may be sufficient and less toxic, we tested a matrix of AA concentrations (1.56, 6.25 and 25 mM) and pretreatment times (5 min, 15 min, 60 min and 24 h) to find an optimal combination. Jurkat cell pretreatment with 1.56 mM AA for just 5 min induced significant ROS mitigation similar to or even better than a 24-hour pretreatment (Figure 3f). This treatment with AA was equally effective when given as a pretreatment, as a post-treatment immediately after light exposure, or when having AA present in the culture media during photoexposure (Figure S3). Furthermore, this short exposure time did not induce cell death compared to untreated cells (Figure 3g). For this reason, cell pretreatment with 1.56 mM AA for 5 min was selected to mitigate ROS in subsequent experiments with gels and other cell types.

### Adding collagen to GelSH precursor before photocrosslinking improved gel porosity

Having established a method to mitigate ROS production, we turned to improving the porosity of the photocrosslinked gel. Given the prevalence of fibrillar structures in the lymph node stroma, and prior reports that addition of fibrillar collagen increased the pore sizes of 3D printed GelMA hydrogel meshes^59^ and porosity of PEG hydrogels,^60^ we hypothesized that addition of collagen would increase both the porosity of the photocrosslinked GelSH gels and the motility of cells encapsulated within them. Therefore, we mixed varied quantities of collagen precursor into the GelSH precursor, photocrosslinked the gels, and allowed an additional (30 min) incubation for thermal gelation. As observed in Figure 2g, the neat GelSH gel had limited porosity when imaged under SEM, while the Col/Fib gel used for comparison was very porous, with visibly interconnected pores. Adding collagen to GelSH precursor prior to photocrosslinking increased gel porosity in a dose-dependent manner (Figure 4), with large pores that qualitatively approached the diameter of those of Col/Fib appearing at 40-60% collagen in the GelSH gel.

**Figure 4.**
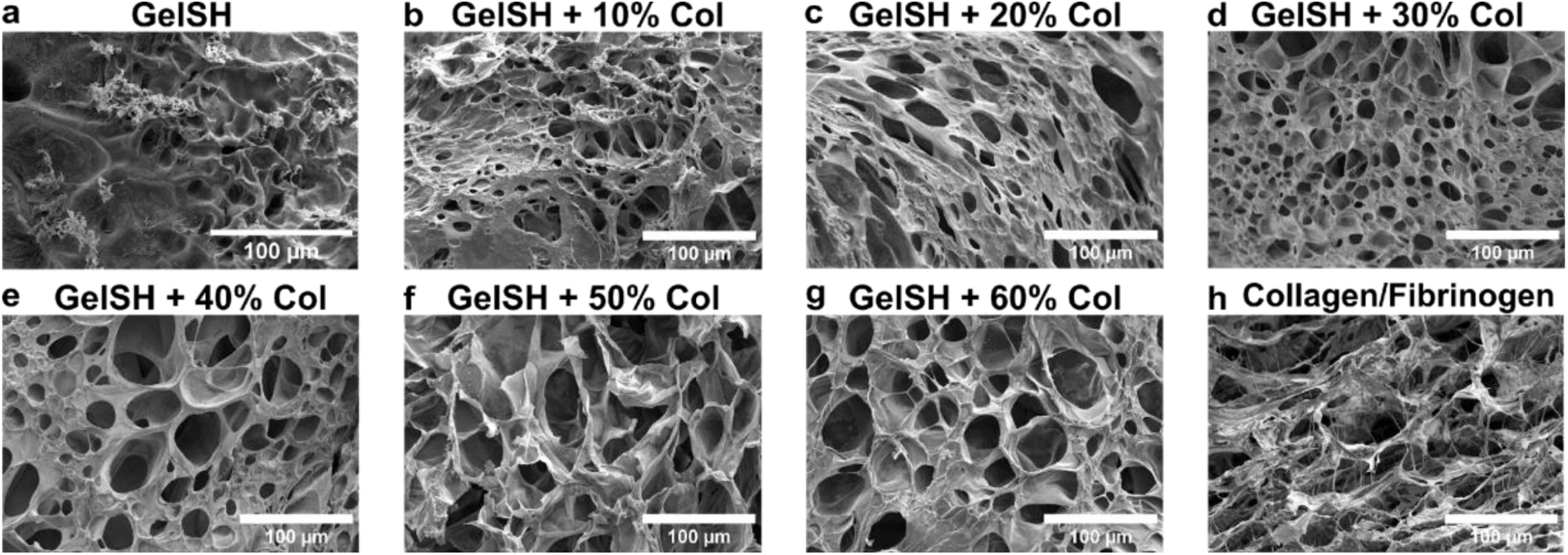
SEM images of GelSH (a), GelSH/collagen mixtures (b-g) and Col/Fib (h) hydrogels, showing differences in gel porosity. Scale bar = 100 μm.

### Increasing GelSH porosity, not quenching ROS, improved multi-day viability of Jurkat cells

Next, we tested the ability of these two approaches, alone or in combination, to preserve viability of Jurkat cells in photocrosslinked 3D GelSH gel cultures. These experiments were conducted in a simple microfluidic chip to allow for high throughput image acquisition, live cell imaging, and spatial photopatterning of multiple cell types in subsequent experiments. Cells were pretreated with 1.56 mM AA for 5 min, resuspended in gel precursor and loaded into the chip for photocrosslinking (GelSH), thermal gelation (Col/Fib) or both (GelSH/collagen gels). The AA pretreatment was washed out prior to gelation, and we did not observe any noticeable impact of the pretreatment on efficacy of gelation. After gelation, media was added to chip reservoirs and the chips were incubated for 3 days. Jurkat cell viability in the neat GelSH gel was poor after 3-day culture (49%), similar to our prior work which showed that viability of primary CD4+ T cells significantly decreased after 24-hour culture in GelSH gels.^10^ Surprisingly, cell pretreatment with AA had no impact on viability, suggesting that the delayed toxicity seen in the gel did not arise from the oxidative insult of ROS production. Instead, increasing the porosity of GelSH hydrogels by adding collagen markedly improved Jurkat cell viability, and viability was greater in GelSH + 60% collagen gels than in GelSH + 40% collagen gels. This result indicated that there was a dose-response relationship, where increased gel porosity translated to improved cell viability. In addition to higher cell viability, Jurkat cells also formed clusters in the GelSH/60% collagen gels (Figure 5c) similar to those observed in the control Col/Fib gels (Figure 5d).

**Figure 5.**
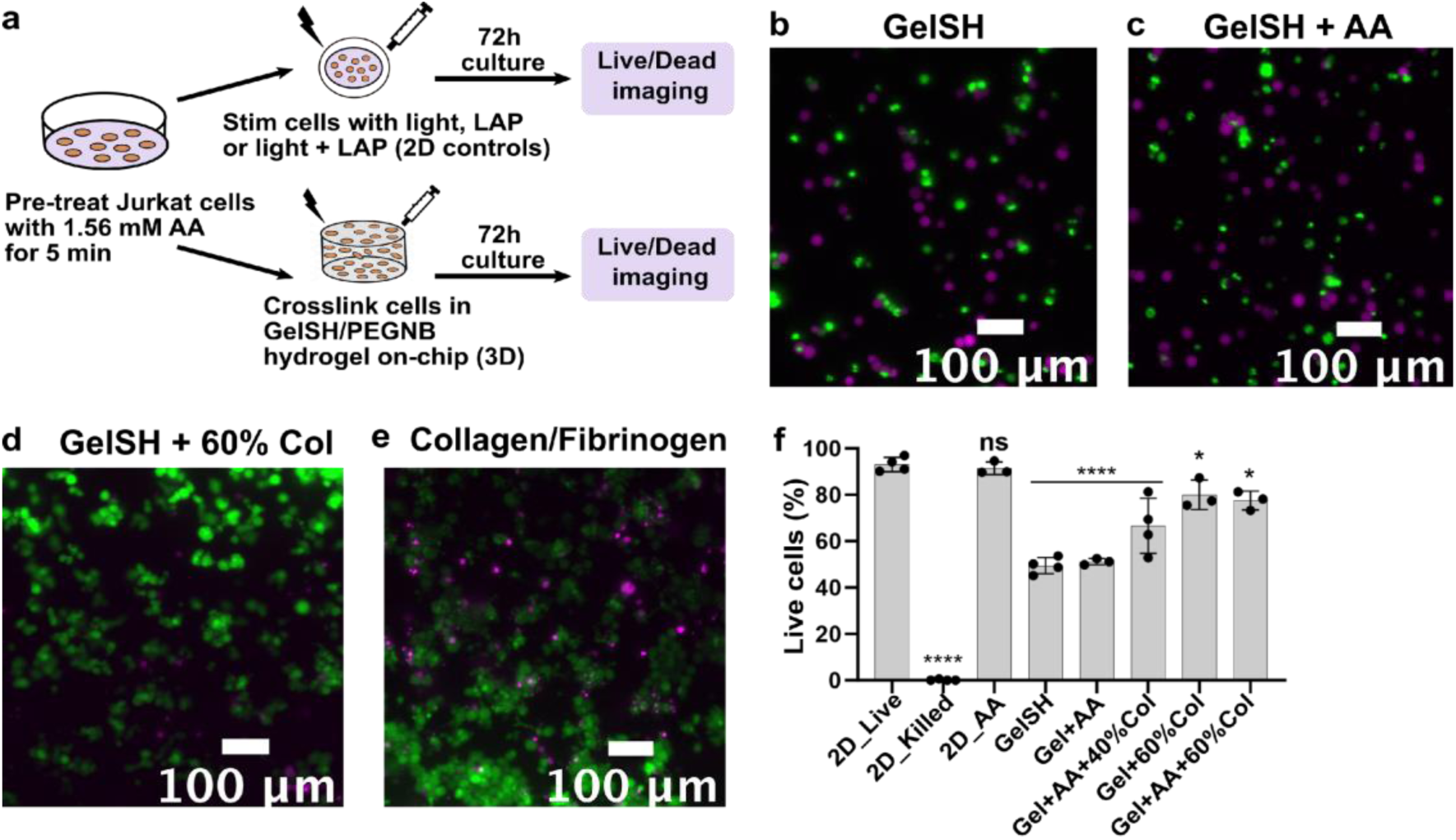
Increasing gel porosity, and not AA pretreatment, improved Jurkat cell viability in photocrosslinked GelSH gels. Representative fluorescence images from viability staining of untreated Jurkat cells in GelSH gel (a), AA-treated cells in GelSH gel (b) and untreated cells in GelSH/60% collagen gel (c) after 72-hour culture. Cells were labeled with calcein AM (green) and DAPI (magenta). Scale bar = 100 μm. (d) Quantification of Jurkat cell viability after 72-hour culture in 2D wells and 3D gels (n = 3). One-way ANOVA with Dunnett’s multiple comparisons, ns p > 0.05, *p < 0.05, ****p < 0.0001 versus untreated cells (2D_Live).

### Fibroblasts spread only in the more porous GelSH + 60% collagen gels despite AA pretreatment

To test the robustness of the above finding, we extended these approaches from the immortal Jurkat cell line to two primary cell types of interest when photopatterning immune cell populations: primary human lymphatic fibroblasts (HLFs) isolated from human lymph nodes, and primary human CD4+ T cells. As with Jurkat cells, HLF exposure to LAP and light + LAP induced a significant increase in ROS levels which was mitigated with AA pretreatment (Figure 6b). Light + LAP induced a 4-fold increase in ROS signal in HLFs, which was less than that in Jurkat cells, which experienced a 16-fold increase. Nevertheless, as with Jurkat cells, the MTS assay indicated that exposure of HLFs to LAP and light + LAP induced significant reduction in metabolic activity (Figure S4a), but not membrane permeability based on results from the LDH assay (Figure S4b). Interestingly, unlike Jurkat cells, the HLFs did not suffer loss of viability in the Live/Dead assay (Figure 6c). After 3-day culture, HLFs remained fully viable in GelSH gel (Figure 6e). However, they were completely unable to spread in GelSH, even with AA pretreatment (Figure 6f). In contrast, HLFs spread in GelSH + 60% collagen gel similar to 2D wells (Figure 6d) and Col/Fib gels (Figure 6h). This trend was retained after 1 week of culture (Figure S5). Similarly, Benton et al^13^ reported limited spreading of aortic valvular interstitial cells in photocrosslinked GelMA hydrogels after 2-day culture. Extensive cell spreading was accomplished only by enzymatic degradation of gel with collagenase, which induced bulk erosion throughout the gel which allowed the cells to spread.^13^ Likewise, Liu and Chan-Park^61^ identified that smooth muscle cells spread better and formed networks in softer photocrosslinked methacrylate-functionalized dextran-gelatin hydrogels whereas spreading was limited in stiffer (more concentrated) gels.

**Figure 6.**
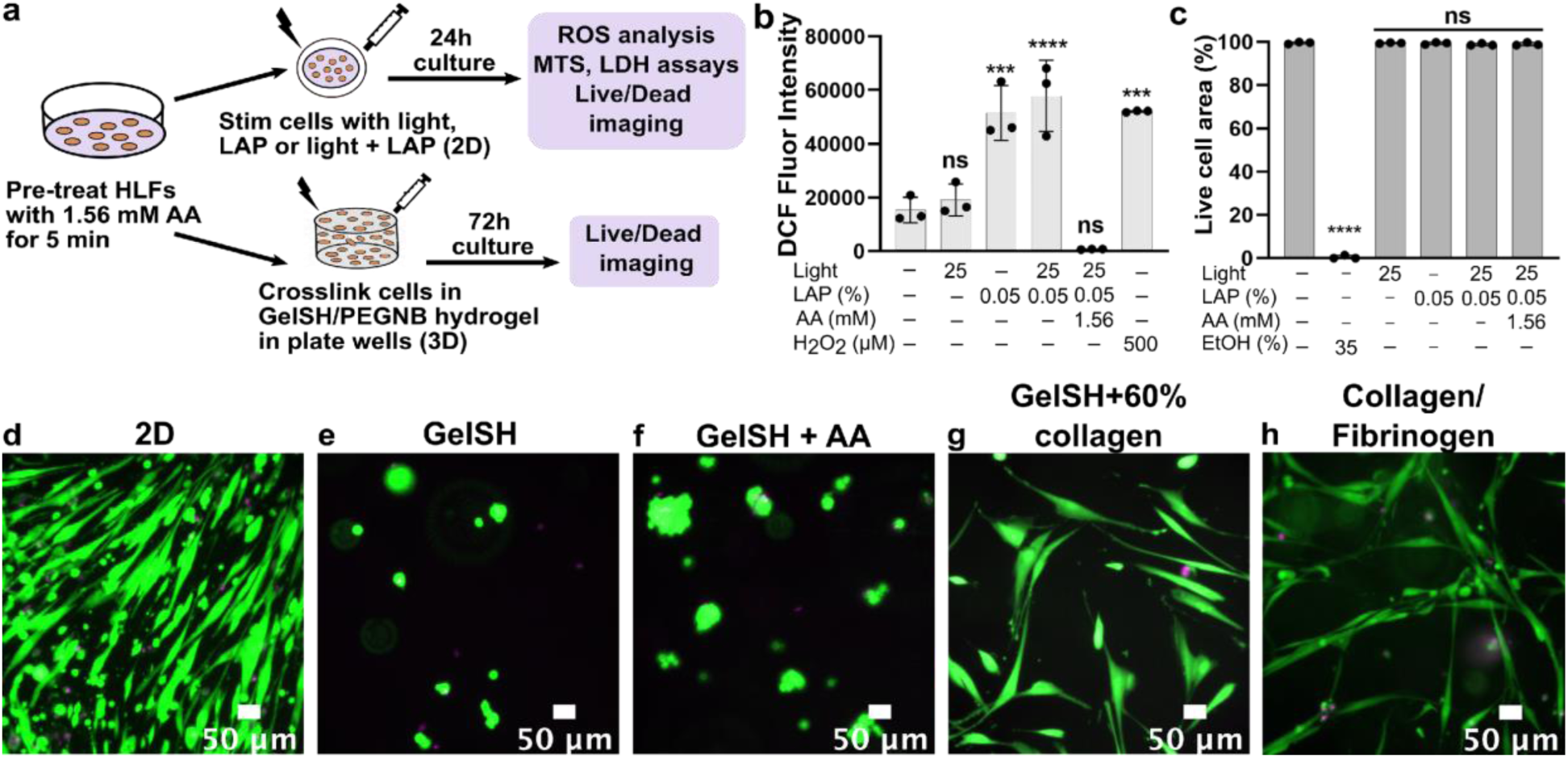
Although AA pretreatment mitigated light + LAP-induced ROS production in HLFs, HLFs spread only in the more porous GelSH + 60% collagen hydrogels. (a) Schematic of experimental setup. (b) DCF fluorescence intensities in HLFs and (c) HLF viability from the Live/Dead assay after 24-hour culture following exposure to light (25 mW/cm^2^), LAP and light + LAP (with or without AA pretreatment) in 2D wells (n = 3). One-way ANOVA with Dunnett’s multiple comparisons, ns p > 0.05, ***p < 0.001, ****p < 0.0001 versus untreated cells. (d-h) Representative fluorescence images of untreated HLFs in 2D wells, untreated HLFs in GelSH, AA-treated HLFs in GelSH, untreated HLFs in GelSH + 60% collagen and untreated HLFs in collagen/fibrinogen gels after 3-day culture and staining with calcein AM (green) and DAPI (magenta). Scale bar = 50 μm.

Remarkably, a critical porosity was also required for HLF spreading in GelSH/collagen gels. As in the neat GelSH gel, HLFs were viable but did not spread in GelSH + 40% collagen gels even after AA pretreatment (Figure S6a), whereas they did spread in the 60% collagen gels (Figure 6c). To test whether the collagen itself was required or simply higher porosity, we compared spreading in GelSH + 60% collagen versus GelSH + 60% PBS gels; the latter was a simple dilution of the precursor with saline. HLFs were able to spread in the GelSH + 60% PBS gels as well as in the GelSH + 60% collagen gel (Figure S6b). This result indicated that HLF spreading in GelSH/collagen gel was driven by increased gel porosity rather than a direct biochemical effect induced by the addition of collagen. We continued to use the collagen, however, as the saline-diluted GelSH was physically weak and challenging to handle.

### Motility of primary human CD4+ T cells improved with increased GelSH porosity and ROS inhibition

Similar to Jurkat cells and HLFs, direct exposure of primary human CD4+ T cells to light and LAP significantly increased ROS, and this effect was mitigated by pretreating cells with AA (Figure 7b). In the absence of gel, photoexposure of CD4+ T cells to light, LAP or the combination thereof had negligible impact on cell viability after 24-hour incubation in 2D wells (Figure 7c).

**Figure 7.**
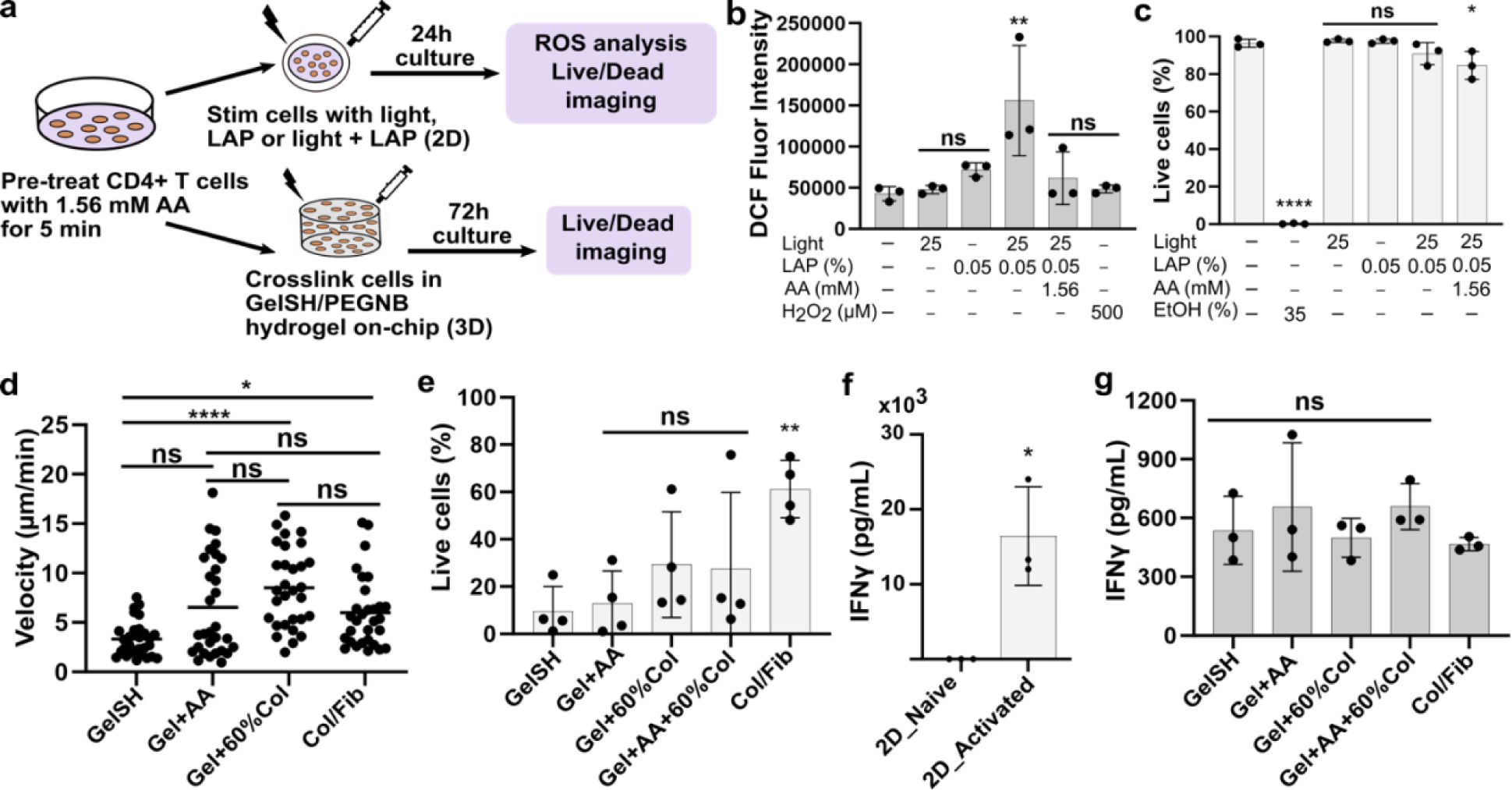
Exposure to light and LAP also increased ROS levels in primary human CD4+ T cells and this effect was mitigated with AA pretreatment. (a) Schematic of experimental setup. (b) DCF fluorescence intensities in primary CD4+ T cells after exposure to light, LAP and light + LAP (with or without AA pretreatment) (n = 3 donors). (c) CD4+ T cell viability from the Live/Dead assay after exposure to light, LAP and light + LAP (with or without AA pretreatment) in 2D wells (n = 3 donors). For b and c, one-way ANOVA with Dunnett’s multiple comparisons, ns p > 0.05, ** p < 0.01, ****p < 0.0001 versus untreated cells. (d) CD4+ T cell motility in hydrogels from live cell imaging after 24-hour culture in media containing CCL21 (1.22 ng/mL). Each dot represents a single cell. Data from one donor is shown here; black line indicates the mean cell velocity. Data were analyzed using a Kruskal-Wallis test with Dunn’s multiple comparisons. **p < 0.01, ****p < 0.0001. Data is representative of n = 3 donors for all conditions. (e) CD4+ T cell viability after 3-day culture in hydrogels (n = 4 donors). Bars show the mean with standard deviation and each dot represents one donor. One-way ANOVA with Dunnett’s multiple comparisons, ns p > 0.05, **p < 0.001 versus GelSH gel. (f, g) IFN**γ** secretion from CD4+ T cells in 2D wells (f) and hydrogels (g) after ImmunoCult™ CD3/CD28 antibody stimulation and 3-day culture (n = 3 donors). One-way ANOVA with Dunnett’s multiple comparisons, ns p > 0.05, *p < 0.01 versus Col/Fib gel.

We tested the effect of mitigating ROS and adding porosity to improve CD4+ T cell motility and multi-day viability. After 24-hour culture in CCL21-containing media, cell tracking analysis revealed that CD4+ T cell velocity in GelSH gels was significantly increased by AA pretreatment, increased porosity and the combination of both factors (Figure 7d, Supplementary Movies S1-S4). Mean cell velocities were 2.8, 10.0, 8.3 and 7.7 µm/min for GelSH, GelSH + AA, GelSH + 60% collagen, GelSH + AA + 60% collagen and Col/Fib groups respectively. The latter three values are comparable to those reported by Stachowiak and Irvine for primary murine CD4+ T cells cultured in CCL21-containing collagen gels.^31^ There was no significant difference in cell motility between the GelSH + AA and GelSH + 60% collagen groups. These results demonstrated that primary CD4+ T cells were sensitive to ROS production and low hydrogel porosity as both factors negatively impacted cell motility in GelSH gels.

Despite this improvement in motility at 24-hr, the viability of CD4+ T cells after 3-days in culture in 3D gels on-chip improved slightly but not significantly in the more porous GelSH + 60% collagen gels (Figure 7e) and was not helped by pretreatment with AA. On the other hand, similar to off-chip 2D controls (Figure 7f), CD4+ T cells in the on-chip gel cultures secreted IFN**γ** upon CD3/CD28 ligation, with no significant differences across experimental groups. (Figure 7g). This result indicates that even though the photoexposure-induced ROS production and the limited porosity of GelSH gels damaged the viability of the cells by three days, they had negligible impact on T cell activation and cytokine secretion. We speculate that cytokine secretion may have occurred early in the culture, prior to loss of viability, though we have not tested that here. Overall, mitigating ROS and increasing hydrogel porosity both improved the motility of primary CD4+ T cells in these photo-crosslinked hydrogels, but only marginally improved cell viability after multi-day culture.

### Porous photo-crosslinkable gels enabled CD4+ T cell migration toward centrally-located HLF cultures on chip

Having identified conditions that increased lymphocyte motility and promoted fibroblast spreading, we tested immune cell co-culture in the photocrosslinked gels. In vivo, T cells are migratory and chemoattracted towards lymphoid fibroblasts. Here, fibroblast-T cell crosstalk was modelled using a multi-lane PDMS microfluidic chip in which primary T cells were loaded in the outer lanes and crosslinked before HLFs were loaded in the center lane and crosslinked, thus proving a spatial organized co-culture (Figure 8a). Consistent with the results from monocultures, HLFs spread in Col/Fib and did not spread at all in the GelSH gel, and T cells remained round and immobile where they were patterned in the GelSH (Figure 8b). In Col/Fib gel, some of the T cells migrated into the center lane, and a small number were observed to co-localize with HLFs there when imaged after 2 days, similar to prior reports.^46,62^

**Figure 8.**
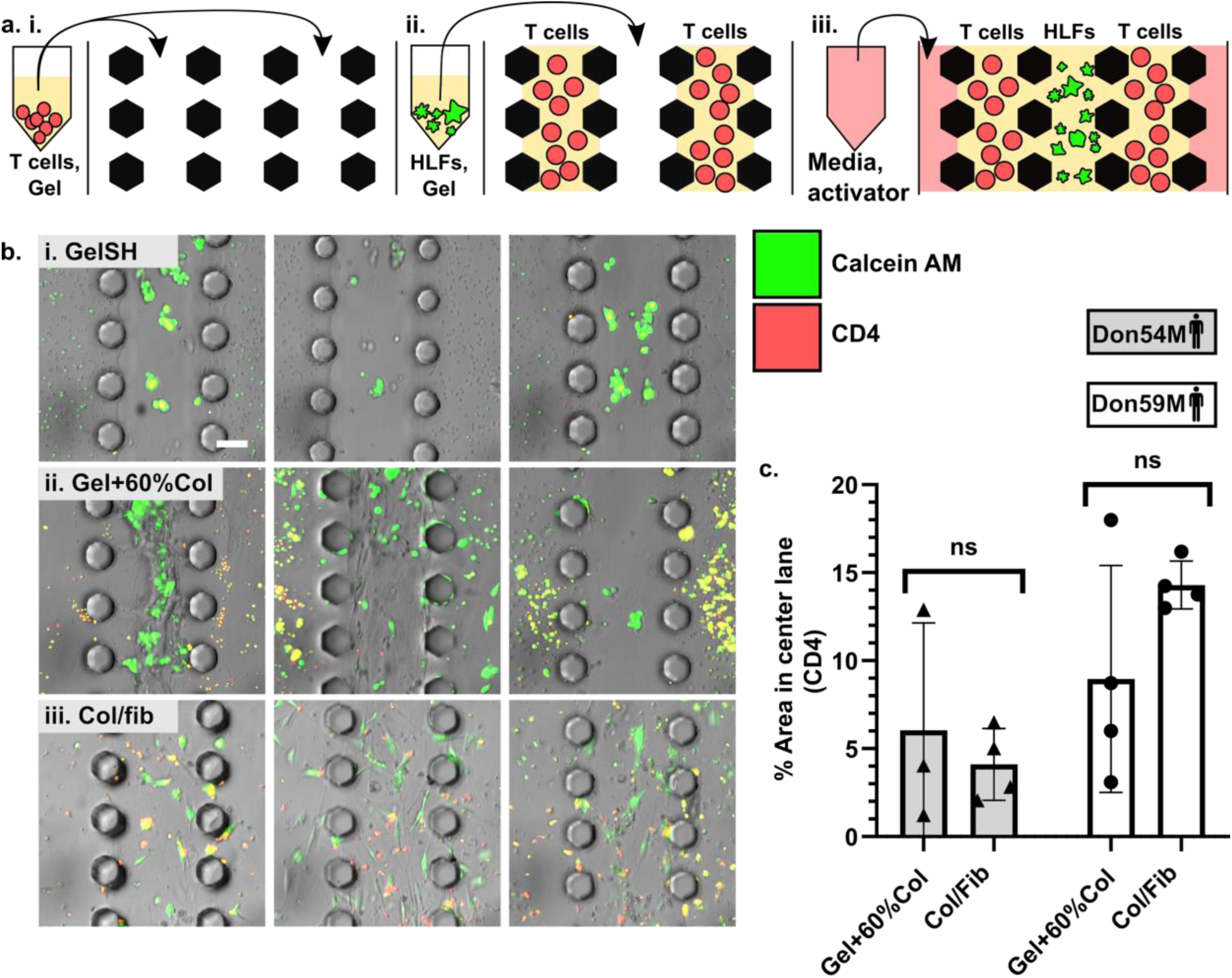
GelSH + 60% collagen hydrogel enabled lymphocyte-stromal colocalization in a microphysiological system. (a) Schematic showing the establishment of CD4+ T cell and HLF co-cultures on a microfluidic chip. (*i*) A T cell-laden gel precursor was loaded, then cured (thermal and/or photo-exposed, depending on the hydrogel formulation). (*ii*) A stromal cell-laden gel precursor was loaded, then cured. (*iii*) Media was added to outer lanes. (b) Representative images of CD4+ T cells and HLFs in each gel after 2-day culture. Within each condition, images were leveled to distinguish cells from background fluorescence signal. Cells were labelled with anti-CD4 (red) and Calcein AM (green). Overlays of brightfield and fluorescence images are shown; three representative images shown for each condition. Scale bar is 100 µm. (c) Quantification of CD4+ T cell area in center lane, for two human donors. Each dot represents one chip (mean of three regions imaged per chip), n = 3 – 4 chips per condition. Data were analyzed using two-way ANOVA with Sidak’s multiple comparisons, ns p > 0.05. Brightfield images showed that no lymphocytes were detected in center lanes for GelSH cultures; these were omitted from the plot due to poor immunofluorescence staining in that gel.

Interestingly, co-cultures in the GelSH + 60% collagen gels yielded HLF spreading and redistribution of the T cells in some regions but not others. Stromal cell morphology in the GelSH + 60% collagen gel was distinct and more variable than the other groups. In two out of six chips, HLFs spread and formed tighter, more contractile networks than were seen in Col/Fib gels. CD4+ T cells appeared to have moved and clustered within their lanes, unlike in neat GelSH, and migrated T cells were frequently visible within the center lane when imaged after 2 days, though with more variability than in Col/Fib gels (Figure 8c). Also, there was less co-localization of T cells with HLFs than was seen in Col/Fib gels. Thus, increasing hydrogel porosity enabled a spatially organized, photo-crosslinked co-coculture that supported T cell migration towards a region containing HLFs, but with regional variability in HLF spreading and T cell migration that remains to be addressed. We hypothesize that the source of the variability may be due to slight segregation between the collagen and GelSH components, as those ingredients have opposite thermal gelation properties. One potential area for future improvement will be to optimize the gel mixing and crosslinking strategies to enable more consistent association between gelatin and collagen fibers.

## CONCLUSION

In conclusion, this work systematically tested the role of ROS generation and porosity in the viability and function of photopatterned, cell-laden hydrogel cultures. The combination of light and LAP used in standard photocrosslinking conditions, even in the absence of polymeric precursors, induced markedly high ROS levels in T cells, both Jurkat and primary, exceeding that of standard peroxide controls. ROS was also induced in HLFs by photocrosslinking, at a level equivalent to that from peroxide. A short, five-minute exposure to low concentrations of AA was sufficient to completely prevent ROS induction in all three types of cells and to maintain viability in 2D culture. This approach to mitigating ROS may have broad utility for other photopatterning strategies, including 3D printing, bioprinting and photolithography-based methods. However, interestingly and unexpectedly, in these 3D cultures the low porosity of the gelatin-based material played a larger role in toxicity than the ROS production. Improving porosity by diluting the gelatin precursor with collagen solution retained the physical stability of the gels, while enabling cell motility and multi-day viability in the photocrosslinked gels. Furthermore, improving hydrogel porosity was also sufficient to support immune cell-cell coculture as demonstrated by T cell migration across culture lanes and into the HLFs lane on the multi-lane chip. While the response was highly variable in GelSH with 60% collagen, there was a clear improvement in lymphocyte motility and fibroblast spreading when compared to GelSH alone. Meanwhile ROS mitigation did not impact gross viability or short-term stimulated IFN-γ secretion, but did improve the motility of primary T cells 24 hours after gelation. We consider it likely that ROS exposure may have other subtle effects on cell differentiation or function. In summary, this work established conditions sufficient for photopatterning of sensitive lymphocytes and lymphatic fibroblasts. While optimization is needed for homogeneity in organized 3D cultures, we anticipate that these strategies will pave the way for photopatterning of these and other cells in organized immunocompetent 3D cultures, bioprinted structures, and organs-on-chip.

## TABLE OF CONTENTS GRAPHIC

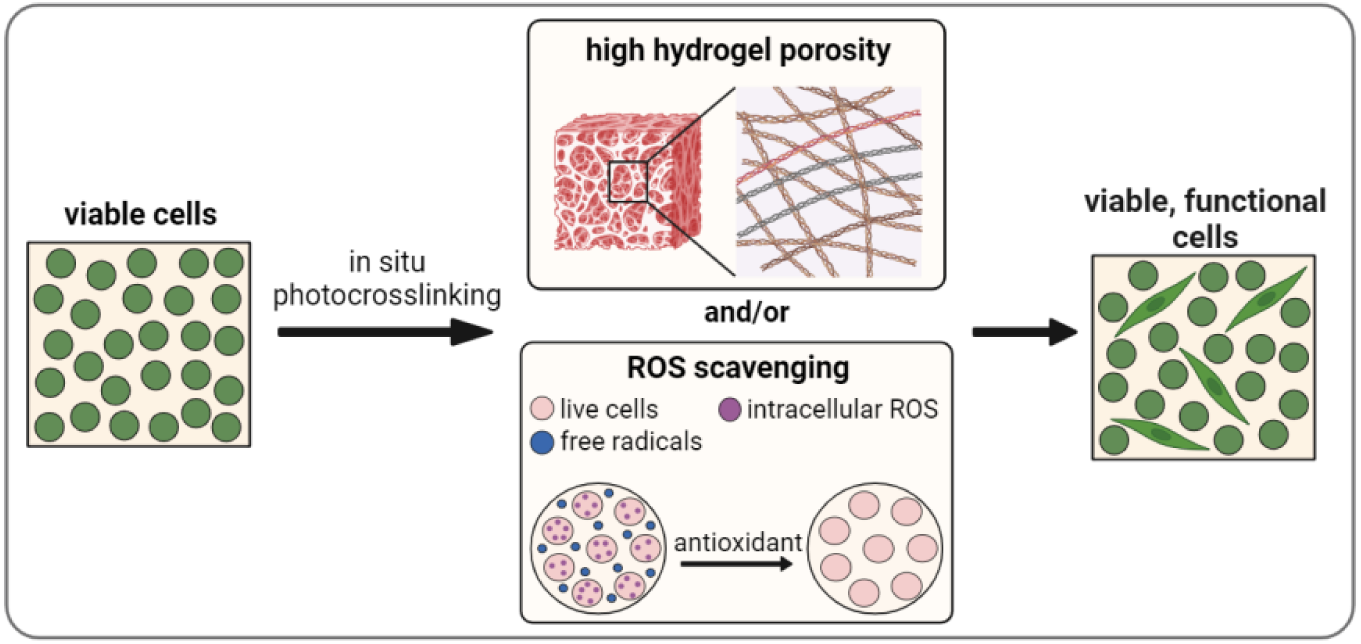

### Author Contributions

TO, JMZ, and RRP led the experimental design, interpreted the results, and drafted the manuscript. TO performed, collected and analyzed data for all experiments. JMZ contributed to experimental set-up, fabrication of microfluidic chips and data collection. AA isolated, characterized, and optimized culture conditions for CD4+ T cells. JHH and JMM provided expertise on cell interactions within biomaterial scaffolds. CJL and TJB provided expertise for design of experiments with T cells and HLFs. TJB passed away in May 2023 and will be greatly missed. All authors reviewed the results and approved the final version of the manuscript.

### Funding Sources

This work was supported by the National Institute of Biomedical Imaging and Bioengineering (NIBIB) under award number U01EB029127 through the National Institutes of Health (NIH), with co-funding from the National Center for Advancing Translational Sciences (NCATS). TO was supported in part by a Summer Research Award through the Global Infectious Diseases Institute at the University of Virginia. JMZ was supported in part by the Graduate Research Fellowship Program through the National Science Foundation. JH was supported by the Virginia Tech Institute for Critical and Applied Science (ICTAS).

## ACKNOWLEDGMENT

Utilization of the Quanta 650 instrument within the University of Virginia’s Nanoscale Materials Characterization Facility (NMCF) was fundamental to this work, and we acknowledge the assistance of Richard White for equipment training. The authors would like to thank the Letteri lab for access to their lyophilizer, and Djuro Raskovic and Isabella Lee for technical assistance during experiments.

## SUPPORTING INFORMATION AVAILABLE

The following supporting information is available free of charge: Supporting figures and methods, captions for supplemental movies.

## REFERENCES

1. Harff, C. & Panoskaltsis-Mortari, A. Tissue engineering of the lymphoid organs. Journal of Immunology and Regenerative Medicine 13, 100049 (2021).

2. Ozulumba, T., Montalbine, A. N., Ortiz-Cárdenas, J. E. & Pompano, R. R. New tools for immunologists: models of lymph node function from cells to tissues. Frontiers in Immunology 14, (2023).

3. Pereira, R. F. & Bártolo, P. J. 3D bioprinting of photocrosslinkable hydrogel constructs. Journal of Applied Polymer Science 132, (2015).

4. Ifkovits, J. L. & Burdick, J. A. Review: photopolymerizable and degradable biomaterials for tissue engineering applications. Tissue Eng 13, 2369–2385 (2007).

5. Rouillard, A. D. et al. Methods for Photocrosslinking Alginate Hydrogel Scaffolds with High Cell Viability. Tissue Engineering Part C: Methods 17, 173–179 (2011).

6. Elisseeff, J. et al. Transdermal photopolymerization for minimally invasive implantation. Proc. Natl. Acad. Sci. U.S.A. 96, 3104–3107 (1999).

7. Chan, V., Zorlutuna, P., Jeong, J. H., Kong, H. & Bashir, R. Three-dimensional photopatterning of hydrogels using stereolithography for long-term cell encapsulation. Lab Chip 10, 2062–2070 (2010).

8. Lim, K. S. et al. Fundamentals and Applications of Photo-Cross-Linking in Bioprinting. Chem. Rev. 120, 10662–10694 (2020).

9. Davey, S. K. et al. Embedded 3D Photopatterning of Hydrogels with Diverse and Complex Architectures for Tissue Engineering and Disease Models. Tissue Eng Part C Methods 21, 1188–1196 (2015).

10. Ortiz-Cárdenas, J. E. et al. Towards spatially-organized organs-on-chip: Photopatterning cell-laden thiol-ene and methacryloyl hydrogels in a microfluidic device. Organs-on-a-Chip 4, 100018 (2022).

11. Mora-Boza, A. et al. Facile Photopatterning of Perfusable Microchannels in Synthetic Hydrogels to Recreate Microphysiological Environments. Adv Mater e2306765 (2023) doi:10.1002/adma.202306765.

12. Greene, T. & Lin, C.-C. Modular Cross-Linking of Gelatin-Based Thiol–Norbornene Hydrogels for in Vitro 3D Culture of Hepatocellular Carcinoma Cells. ACS Biomater. Sci. Eng. 1, 1314–1323 (2015).

13. Benton, J. A., DeForest, C. A., Vivekanandan, V. & Anseth, K. S. Photocrosslinking of Gelatin Macromers to Synthesize Porous Hydrogels That Promote Valvular Interstitial Cell Function. Tissue Engineering Part A 15, 3221–3230 (2009).

14. Bhise, N. S. et al. A liver-on-a-chip platform with bioprinted hepatic spheroids. Biofabrication 8, 014101 (2016).

15. R Ibañez, R. I., et al. 3D-Printed Gelatin Methacrylate Scaffolds with Controlled Architecture and Stiffness Modulate the Fibroblast Phenotype towards Dermal Regeneration. Polymers (Basel) 13, 2510 (2021).

16. Jabbari, E., Sarvestani, S. K., Daneshian, L. & Moeinzadeh, S. Optimum 3D Matrix Stiffness for Maintenance of Cancer Stem Cells Is Dependent on Tissue Origin of Cancer Cells. PLOS ONE 10, e0132377 (2015).

17. Fan, L. et al. Directing Induced Pluripotent Stem Cell Derived Neural Stem Cell Fate with a Three-Dimensional Biomimetic Hydrogel for Spinal Cord Injury Repair. ACS Appl. Mater. Interfaces 10, 17742–17755 (2018).

18. Brown, A. et al. Engineering PEG-based hydrogels to foster efficient endothelial network formation in free-swelling and confined microenvironments. Biomaterials 243, 119921 (2020).

19. Kappes, U. P., Luo, D., Potter, M., Schulmeister, K. & Rünger, T. M. Short– and long-wave UV light (UVB and UVA) induce similar mutations in human skin cells. J Invest Dermatol 126, 667–675 (2006).

20. Klak, M. et al. Irradiation with 365 nm and 405 nm wavelength shows differences in DNA damage of swine pancreatic islets. PLoS One 15, e0235052 (2020).

21. Billiet, T., Gevaert, E., De Schryver, T., Cornelissen, M. & Dubruel, P. The 3D printing of gelatin methacrylamide cell-laden tissue-engineered constructs with high cell viability. Biomaterials 35, 49–62 (2014).

22. Lin, H. et al. Application of visible light-based projection stereolithography for live cell-scaffold fabrication with designed architecture. Biomaterials 34, 331–339 (2013).

23. Bartnikowski, M., Bartnikowski, N. J., Woodruff, M. A., Schrobback, K. & Klein, T. J. Protective effects of reactive functional groups on chondrocytes in photocrosslinkable hydrogel systems. Acta Biomater 27, 66–76 (2015).

24. Williams, C. G., Malik, A. N., Kim, T. K., Manson, P. N. & Elisseeff, J. H. Variable cytocompatibility of six cell lines with photoinitiators used for polymerizing hydrogels and cell encapsulation. Biomaterials 26, 1211–1218 (2005).

25. Yeh, J. et al. Micromolding of shape-controlled, harvestable cell-laden hydrogels. Biomaterials 27, 5391–5398 (2006).

26. Jiang, Z., Jiang, K., McBride, R. & Oakey, J. S. Comparative cytocompatibility of multiple candidate cell types to photoencapsulation in PEGNB/PEGDA macroscale or microscale hydrogels. Biomed. Mater. 13, 065012 (2018).

27. Xu, H., Casillas, J., Krishnamoorthy, S. & Xu, C. Effects of Irgacure 2959 and lithium phenyl-2,4,6-trimethylbenzoylphosphinate on cell viability, physical properties, and microstructure in 3D bioprinting of vascular-like constructs. Biomedical Materials 15, 055021 (2020).

28. Nguyen, A. K., Goering, P. L., Elespuru, R. K., Sarkar Das, S. & Narayan, R. J. The Photoinitiator Lithium Phenyl (2,4,6-Trimethylbenzoyl) Phosphinate with Exposure to 405 nm Light Is Cytotoxic to Mammalian Cells but Not Mutagenic in Bacterial Reverse Mutation Assays. Polymers (Basel) 12, 1489 (2020).

29. Roberts, J. J. & Bryant, S. J. Comparison of Photopolymerizable Thiol-ene PEG and Acrylate-Based PEG Hydrogels for Cartilage Development. Biomaterials 34, 9969–9979 (2013).

30. Mũnoz, Z., Shih, H. & Lin, C.-C. Gelatin hydrogels formed by orthogonal thiol– norbornene photochemistry for cell encapsulation. Biomater. Sci. 2, 1063–1072 (2014).

31. Stachowiak, A. N. & Irvine, D. J. Inverse opal hydrogel-collagen composite scaffolds as a supportive microenvironment for immune cell migration. J. Biomed. Mater. Res. 85A, 815–828 (2008).

32. Choi, D. J. et al. Effect of the pore size in a 3D bioprinted gelatin scaffold on fibroblast proliferation. Journal of Industrial and Engineering Chemistry 67, 388–395 (2018).

33. Boonrungsiman, S. et al. Shape and surface properties of titanate nanomaterials influence differential cellular uptake behavior and biological responses in THP-1 cells. Biochem Biophys Rep 9, 203–210 (2017).

34. Kalyanaraman, B. et al. Measuring reactive oxygen and nitrogen species with fluorescent probes: challenges and limitations. Free Radic Biol Med 52, 1–6 (2012).

35. Cory, A. H., Owen, T. C., Barltrop, J. A. & Cory, J. G. Use of an Aqueous Soluble Tetrazolium/Formazan Assay for Cell Growth Assays in Culture. Cancer Communications 3, 207–212 (1991).

36. Abe, K. & Matsuki, N. Measurement of cellular 3-(4,5-dimethylthiazol-2-yl)-2,5-diphenyltetrazolium bromide (MTT) reduction activity and lactate dehydrogenase release using MTT. Neuroscience Research 38, 325–329 (2000).

37. Collagen Type I, Rat Tail or Bovine | 3D Cell Culture Gels & Coating. Ibidi https://ibidi.com/cell-culture-microscopy/107-collagen-type-i.html.

38. Piccinini, F., Kiss, A. & Horvath, P. CellTracker (not only) for dummies. Bioinformatics 32, 955–957 (2016).

39. Fairbanks, B. D., Schwartz, M. P., Bowman, C. N. & Anseth, K. S. Photoinitiated polymerization of PEG-diacrylate with lithium phenyl-2,4,6-trimethylbenzoylphosphinate: polymerization rate and cytocompatibility. Biomaterials 30, 6702–6707 (2009).

40. Li, W.-G. et al. H2O2-induced O2⋅‒ Production by a Non-phagocytic NAD(P)H Oxidase Causes Oxidant Injury. J Biol Chem 276, 29251–29256 (2001).

41. Wijeratne, S. S. K., Cuppett, S. L. & Schlegel, V. Hydrogen peroxide induced oxidative stress damage and antioxidant enzyme response in Caco-2 human colon cells. J Agric Food Chem 53, 8768–8774 (2005).

42. Coyle, C. H. & Kader, K. N. Mechanisms of H2O2-induced oxidative stress in endothelial cells exposed to physiologic shear stress. ASAIO J 53, 17–22 (2007).

43. Wang, X. & Roper, M. G. Measurement of DCF fluorescence as a measure of reactive oxygen species in murine islets of Langerhans. Anal Methods 6, 3019–3024 (2014).

44. Ruskowitz, E. R. & DeForest, C. A. Proteome-wide Analysis of Cellular Response to Ultraviolet Light for Biomaterial Synthesis and Modification. ACS Biomater. Sci. Eng. 5, 2111–2116 (2019).

45. Choi, D. J. et al. Effect of the pore size in a 3D bioprinted gelatin scaffold on fibroblast proliferation. Journal of Industrial and Engineering Chemistry 67, 388–395 (2018).

46. Kim, J. et al. Characterizing natural hydrogel for reconstruction of three-dimensional lymphoid stromal network to model T-cell interactions. J Biomed Mater Res A 103, 2701–2710 (2015).

47. Munson, J. M., Bellamkonda, R. V. & Swartz, M. A. Interstitial Flow in a 3D Microenvironment Increases Glioma Invasion by a CXCR4-Dependent Mechanism. Cancer Res 73, 1536–1546 (2013).

48. Humayun, M. et al. Elucidating cancer-vascular paracrine signaling using a human organotypic breast cancer cell extravasation model. Biomaterials 270, 120640 (2021).

49. Caliari, S. R. & Burdick, J. A. A practical guide to hydrogels for cell culture. Nature Methods 13, 405–414 (2016).

50. Sadjadi, Z., Zhao, R., Hoth, M., Qu, B. & Rieger, H. Migration of Cytotoxic T Lymphocytes in 3D Collagen Matrices. Biophysical Journal 119, 2141–2152 (2020).

51. van Steen, A. C. I. et al. Transendothelial migration induces differential migration dynamics of leukocytes in tissue matrix. J Cell Sci 134, jcs258690 (2021).

52. Zhitkovich, A. N-Acetylcysteine: Antioxidant, Aldehyde Scavenger, and More. Chem. Res. Toxicol. 32, 1318–1319 (2019).

53. Mlejnek, P. Direct Interaction between N-Acetylcysteine and Cytotoxic Electrophile—An Overlooked In Vitro Mechanism of Protection. Antioxidants (Basel) 11, 1485 (2022).

54. Millea, P. J. N-acetylcysteine: multiple clinical applications. Am Fam Physician 80, 265– 269 (2009).

55. Carr, A. C. & Frei, B. Toward a new recommended dietary allowance for vitamin C based on antioxidant and health effects in humans2. The American Journal of Clinical Nutrition 69, 1086–1107 (1999).

56. Njus, D., Kelley, P. M., Tu, Y.-J. & Schlegel, H. B. Ascorbic acid: The chemistry underlying its antioxidant properties. Free Radic Biol Med 159, 37–43 (2020).

57. Yimcharoen, M. et al. Effects of ascorbic acid supplementation on oxidative stress markers in healthy women following a single bout of exercise. J Int Soc Sports Nutr 16, 2 (2019).

58. Kesarwani, P., Murali, A. K., Al-Khami, A. A. & Mehrotra, S. Redox Regulation of T-Cell Function: From Molecular Mechanisms to Significance in Human Health and Disease. Antioxidants & Redox Signaling 18, 1497–1534 (2013).

59. Nagaraj, A., Etxeberria, A. E., Naffa, R., Zidan, G. & Seyfoddin, A. 3D-Printed Hybrid Collagen/GelMA Hydrogels for Tissue Engineering Applications. Biology (Basel) 11, 1561 (2022).

60. Chen, J.-X. et al. Fabrication of tough poly(ethylene glycol)/collagen double network hydrogels for tissue engineering. Journal of Biomedical Materials Research Part A 106, 192– 200 (2018).

61. Liu, Y. & Chan-Park, M. B. A biomimetic hydrogel based on methacrylated dextran-graft-lysine and gelatin for 3D smooth muscle cell culture. Biomaterials 31, 1158–1170 (2010).

62. Brian J. Kwee, Adovi Akue, & Kyung E. Sung. On-chip human lymph node stromal network for evaluating dendritic cell and T-cell trafficking. bioRxiv 2023.03.21.533042 (2023) doi:10.1101/2023.03.21.533042.

