## Supplementary Information for "Mitigating reactive oxygen species production and increasing gel porosity improves lymphocyte motility and fibroblast spreading in photocrosslinked gelatin-thiol hydrogels"

Electronic Supplementary Information for

**Contents**:

- Supplemental Figure S1: Jurkat cell viability after photoexposure and subsequent 24-hour culture
- Supplemental Figure S2: Antioxidant mitigation of $H_2O_2$-induced ROS in Jurkat cells
- Supplemental Figure S3: Sequence of AA pretreatment had no effect on efficacy of ROS mitigation in Jurkat cells
- Supplemental Figure S4: HLF viability after photoexposure and subsequent 24-hour culture
- Supplemental Figure S5: Fluorescence images of HLFs after 1-week culture
- Supplemental Figure S6: Fluorescence images of HLFs after 3-day culture in GelSH + 40% collagen and GelSH + 60% PBS gels
- Supplemental Figure S7: Viability of activated CD4 T cells in 2D wells
- Supplemental Protocol S1: Microscopy for brightfield and fluorescence images
- Supplemental Movies S1 – S4

**Supplementary Figures**

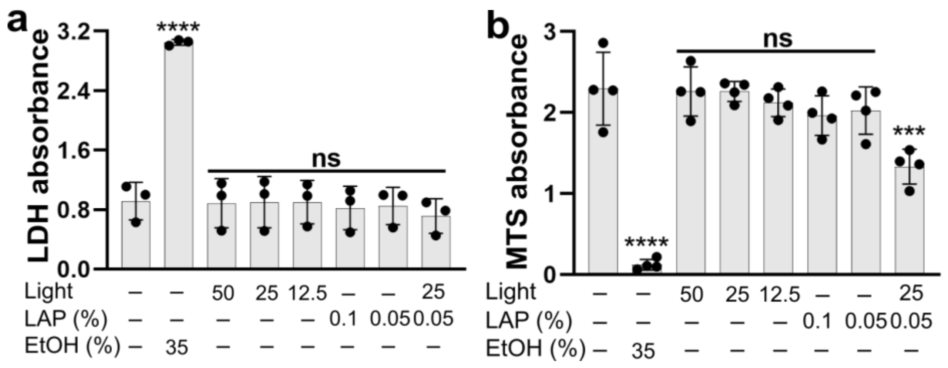


**Figure S1.** Jurkat cell viability after photoexposure and subsequent 24-hour culture. (a) LDH absorbance signal in Jurkat cells 24 h after exposure to light, LAP and light + LAP (n = 3). Ethanol was used as a positive control for cell damage. (b) MTS absorbance signal in Jurkat cells 24 h after exposure to light, LAP and light + LAP (n = 4). One-way ANOVA with Dunnett's multiple comparisons, ns $p > 0.05$, ****$p < 0.0001$ versus untreated cells.

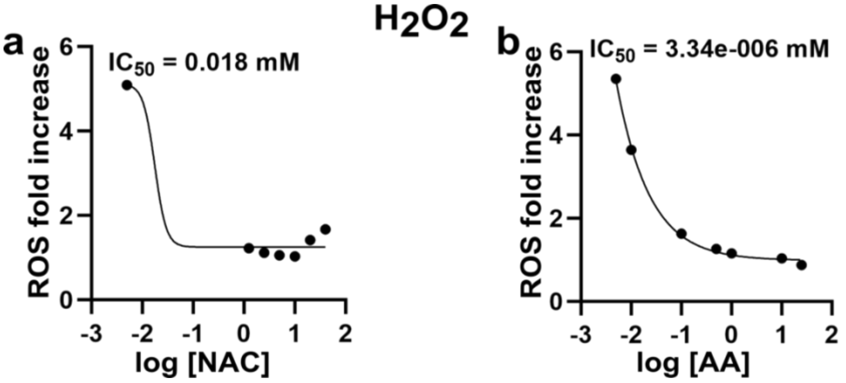


**Figure S2.** Antioxidant mitigation of $H_2O_2$-induced ROS in Jurkat cells. ROS fold increase in Jurkat cells after overnight antioxidant pretreatment with NAC (1.25 – 40 mM) (a) or AA (1.56 – 25 mM) (b) and subsequent 24h exposure to 500 µM $H_2O_2$ (n = 3). Values were fit with a 4-parameter log(inhibitor) vs response curve (*Y=Bottom + (Top-Bottom)/(1+10^((LogIC50-X)*HillSlope)))*.

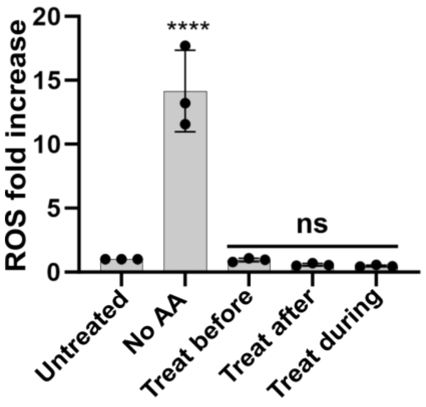


**Figure S3.** Sequence of AA pretreatment had no effect on efficacy of ROS mitigation in Jurkat cells. Jurkat cells were exposed to light (25 mW/cm$^2$) + 0.05% LAP in 2D wells before, during, or after treatment with 1.56 mM AA, incubated for 24 h after which ROS levels were measured. Pre- and post-treatments with AA were conducted for 5 min prior to rinsing it out. One-way ANOVA with Dunnett's multiple comparisons, ns $p > 0.05$, ****$p < 0.0001$ versus untreated cells.

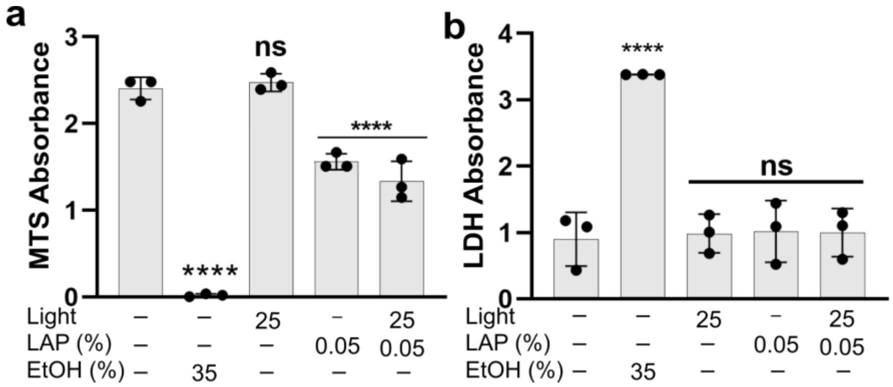


**Figure S4.** Human lymphatic fibroblast (HLF) viability after photoexposure and subsequent 24-hour culture. MTS (a) and LDH (b) absorbance signals in HLFs 24 h after exposure to light, LAP and light + LAP ($n = 3$). One-way ANOVA with Dunnett's multiple comparisons, ns $p > 0.05$, ****$p < 0.0001$ versus untreated cells.

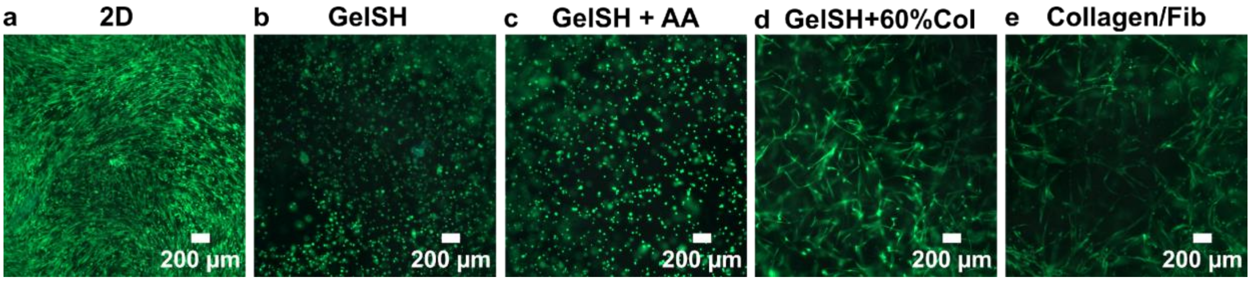


**Figure S5.** Fluorescence images of untreated HLFs in 2D wells (a), untreated HLFs in GelSH (b), AA-treated HLFs in GelSH (c), and untreated HLFs in GelSH + 60% collagen (d) and collagen/fibrinogen (h) gels after 1-week culture and staining with calcein AM (green) and DAPI (blue). Scale bar = 200 µm.

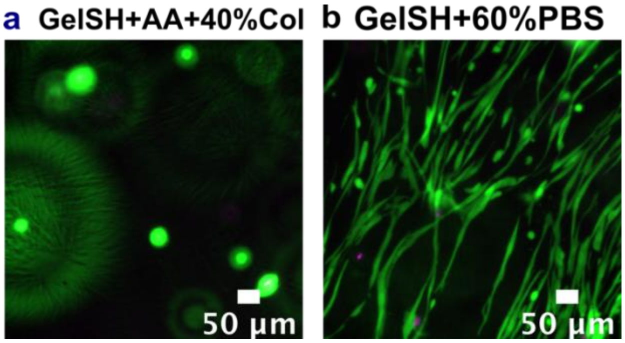


**Figure S6.** Fluorescence images of AA-treated HLFs in GelSH + 40% collagen (a) and untreated HLFs in GelSH + 60% PBS (b) after 3-day culture and staining with calcein AM (green) and DAPI (magenta). Scale bar = 50 µm.

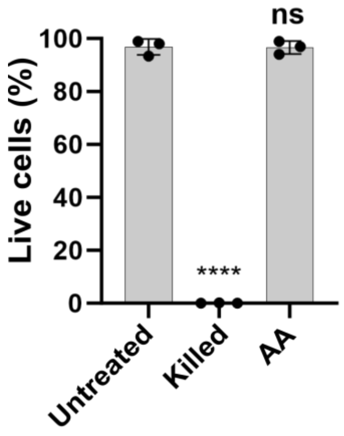


**Figure S7.** Viability of activated CD4 T cells from the Live/Dead assay after 3-day culture in 2D wells. For the AA group, cells were treated for 5 min followed by 3-day culture. One-way ANOVA with Dunnett's multiple comparisons, ns $p > 0.05$, ****$p < 0.0001$ versus untreated cells.

### Protocols

#### Protocol S1: Microscopy for brightfield and fluorescence images.

Images were acquired on a Zeiss AxioZoom widefield microscope equipped with an HXP 200C metal halide lamp, PlanNeoFluor Z 1× objective (0.25 NA, FWD 56 mm), and Axiocam 506 mono camera. Brightfield images were acquired using transmitted light. Filter cubes for fluorescence imaging included DAPI (Ex: 365, Em: 445/50, Zeiss Filter Set, #49), EGFP (Ex: 470/40, Em: 525/50, #38), Rhodamine (Ex: 550/25, Em: 605/70, #43) and Cy5 (Ex: 587/25, Em: 647/70, #64). Images were collected using Zen Blue software (v3.4), and analyzed in ImageJ (v1.53t).

### Movies

For all movies, brightfield images were acquired using transmitted light in time series mode (1 image was collected every 30 sec for 10 cycles). Individual cells (15 – 30) were tracked for image analysis using Cell Tracker (v1.1).

**Movie S1:** Motility of untreated CD4+ T cells in GelSH hydrogel

**Movie S2:** Motility of ascorbic acid treated CD4+ T cells in GelSH hydrogel

**Movie S3:** Motility of CD4+ T cells in GelSH + 60% collagen hydrogel

**Movie S4:** Motility of untreated CD4+ T cells in collagen/fibrinogen hydrogel
